## Supporting Figures and Table for "Deciphering the alphabet of disorder — Glu and Asp act differently on local but not global properties"

SUPPORTING TABLES

TABLE S1: Sequences of *S. pombe* Dss1 wildtype and Glu/Asp variants

SUPPORTING FIGURES

FIGURE S1: MSA logo for the Dss1_Sem1 family

FIGURE S2: Influence of background dataset on composition profile

FIGURE S3: Enrichment profiles of high-resolution X-ray crystal structures

FIGURE S4: NMR titrations of Dss1 variants with ubiquitin

FIGURE S5: Comparing experimental and theoretical SAXS profiles

FIGURE S6: Helix formation time series, peptides of Dss1

FIGURE S7: Helix formation time series, full length Dss1

FIGURE S8: Helix population, simulations

FIGURE S9: Salt bridge frequency, simulations

#### Table S1: Sequences of *S. pombe* Dss1 wildtype and Glu/Asp variants

| | |
|---|---|
| **Dss1 WT N71C** | MSRAALPSLE NLEDDDEFED FATENWPMKD TELDTGDDTL WENNWDDEDI GDDDFSVQLQ AELKKKGVAA C |
| **Dss1 all E N71C** | MSRAALPSLE NLEEEEEFEE FATENWPMKE TELETGEETL WENNWEEEEI GEEEFSVQLQ AELKKKGVAA C |
| **Dss1 all D N71C** | MSRAALPSLD NLDDDDDFDD FATDNWPMKD TDLDTGDDTL WDNNWDDDDI GDDDFSVQLQ ADLKKKGVAA C |
| **Dss1 swap N71C** | MSRAALPSLD NLDEEEDFDE FATDNWPMKE TDLETGEETL WDNNWEEDEI GEEEFSVQLQ ADLKKKGVAA C |

#### Figure S1. MSA logo for Dss1/Sem1 family

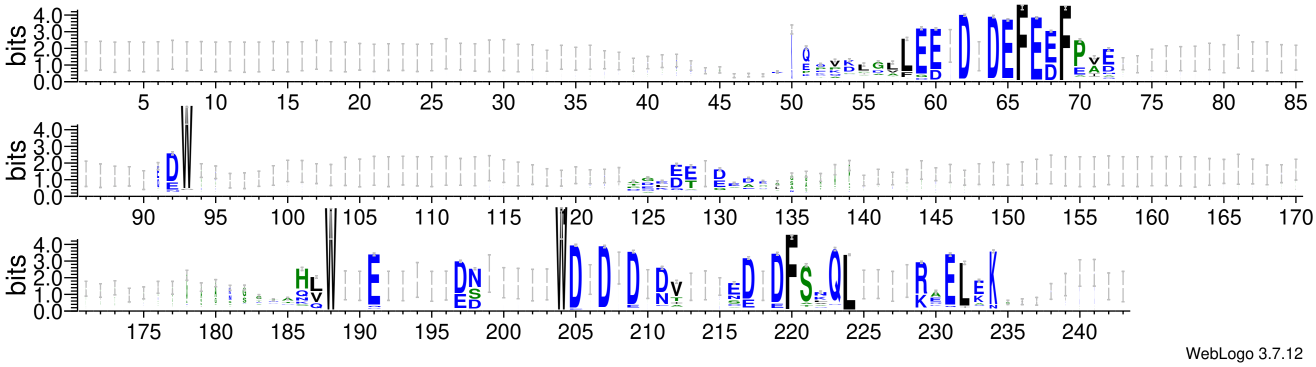


Figure S1: Graphical representation of multiple sequence alignment of all 1612 sequences in the Pfam Dss1/Sem1 family (PF05160), with information content on the y-axis. The graphic was made using WebLogo.

Figure S2: Influence of background dataset on composition profile

A

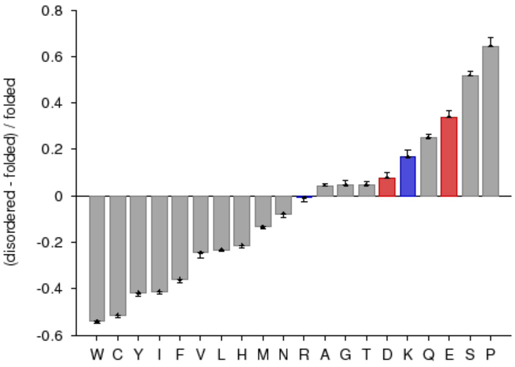


B

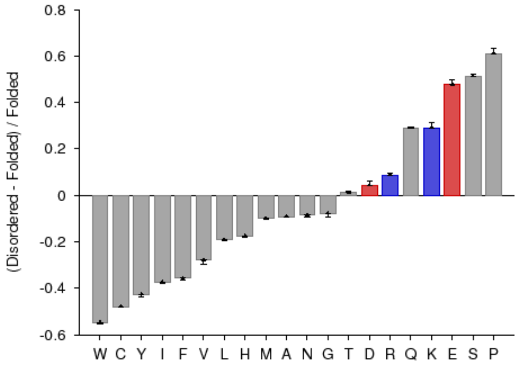


C

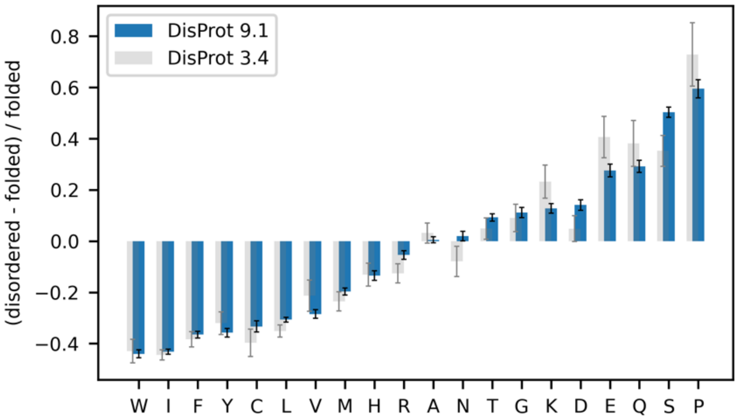


Figure S2: Computed fractional difference between a query set consisting of all non-ambiguous sequences in DisProt v. 8.1 and either: (A) a background set consisting of sequences from PDBselect25 (standard Composition Profiler background set), (B) a background set consisting of sequences from selected proteins with high resolution crystal structures (http://kinemage.biochem.duke.edu/databases/top500.php). Positively charged residues are shown in blue and negatively charged residues are shown in red. (C) Computed fractional difference between a query set consisting of all non-ambiguous sequences in DisProt v. 9.1 or v. 3.4 using a broadly defined background including proteins with low resolution. The fractional difference is an average over 10,000 bootstrap iterations.

### Figure S3: Composition profile of high-resolution X-ray crystal structures

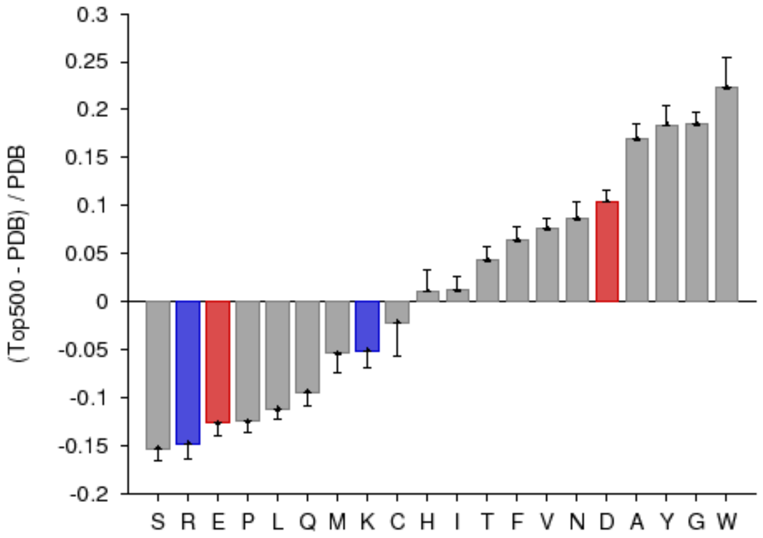


Figure S3: Composition profile of a selection of high-resolution X-ray crystal structures (http://kinemage.biochem.duke.edu/databases/top500.php) with a set of folded proteins from UniProtKB as background. The fractional difference is an average over 10,000 bootstrap iterations. Positively charged residues are shown in blue and negatively charged residues are shown in red.

Figure S4: NMR titrations of Dss1 variants with ubiquitin

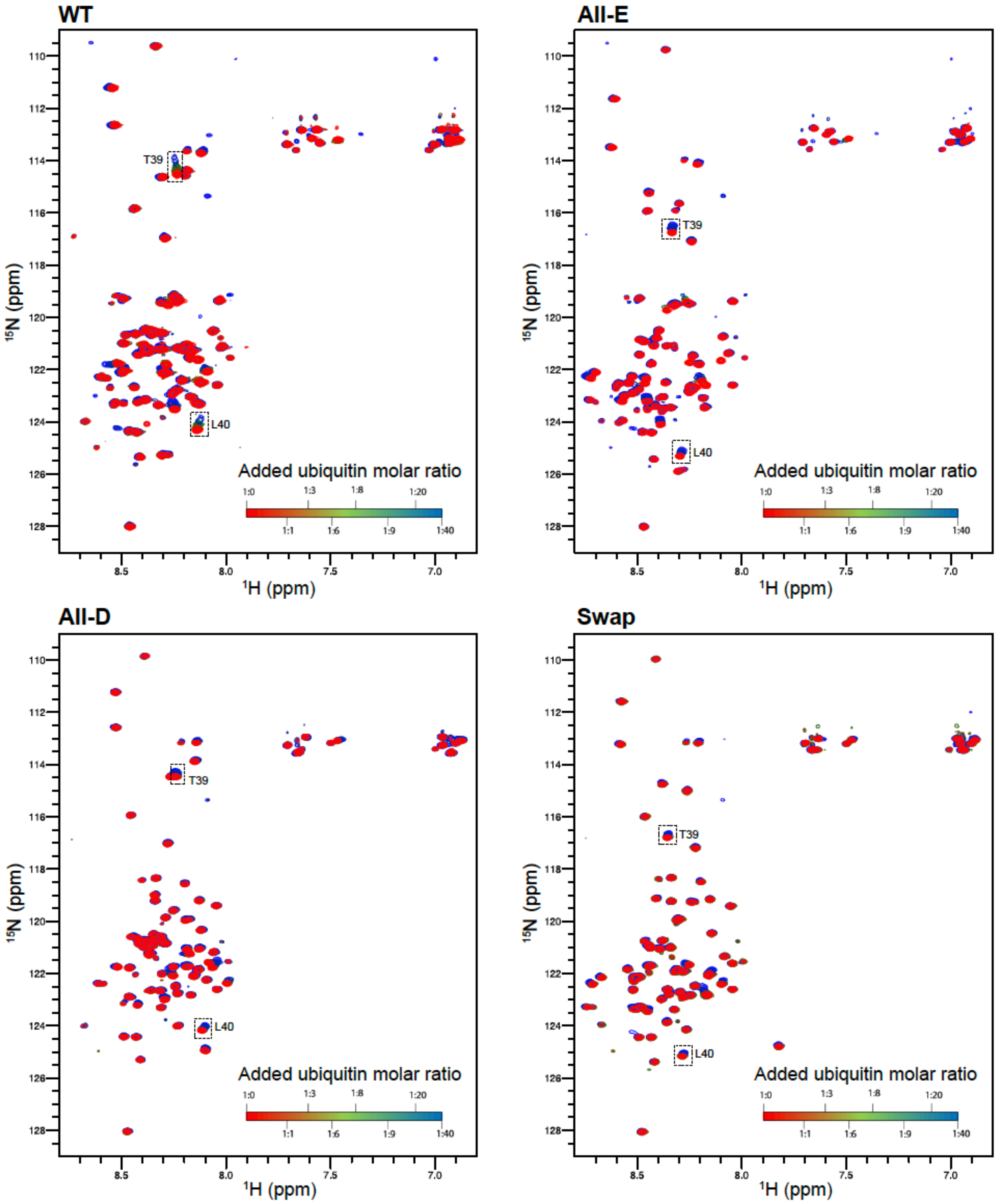


Figure *S4*: NMR titrations of 15N-labeled Dss1 variants All-E, All-D and Swap with mono-ubiquitin to 20 times molar excess

Figure S5: Comparing experimental and theoretical SAXS profiles

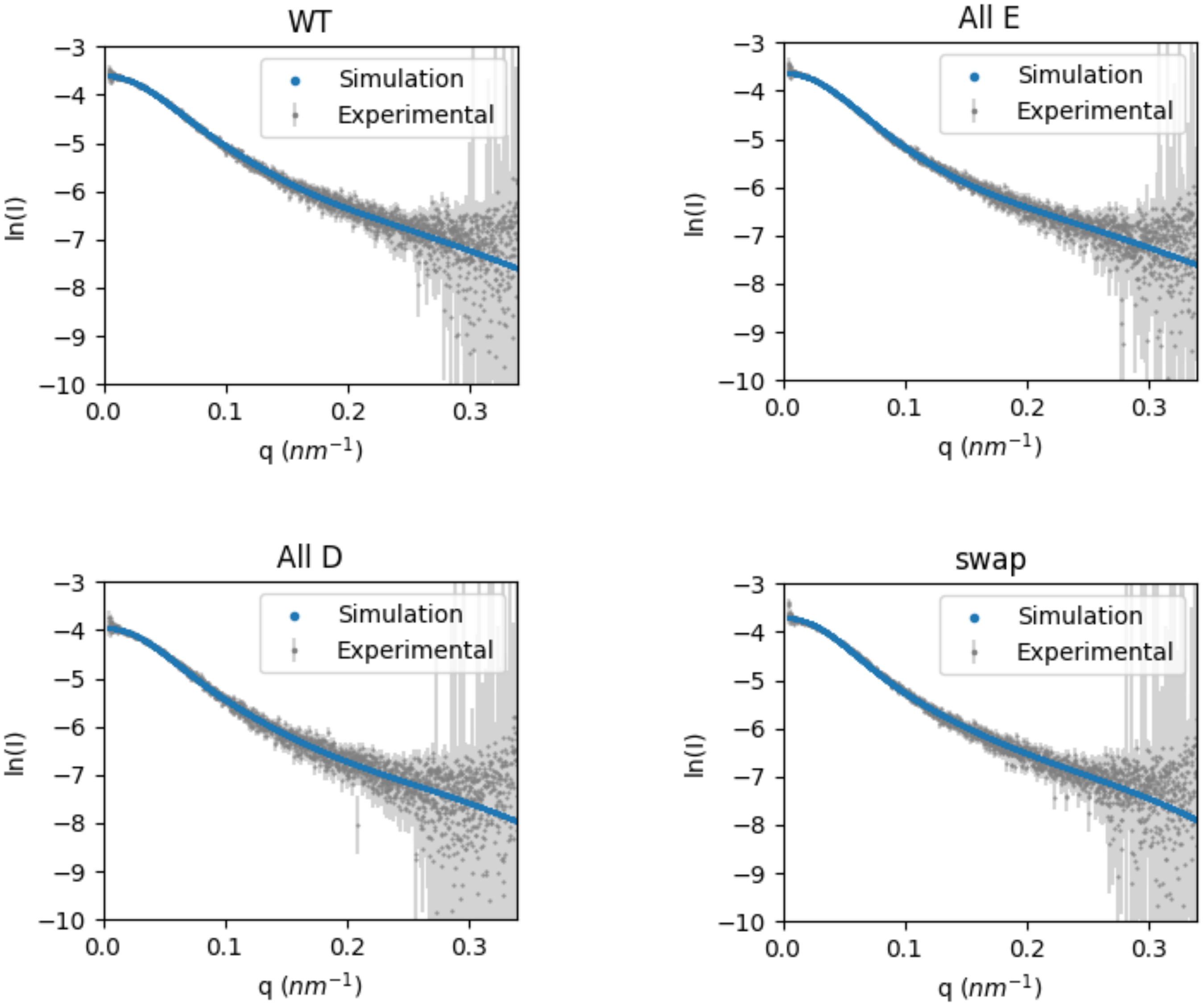


Figure S5: Theoretical SAXS profiles calculated from the MD simulations and the experimental SAXS profiles of Dss1 wildtype and variants. Errors on the experimental intensities are corrected by a Bayesian Indirect Fourier transformation (BIFT) correction factor.

### Figure S6: Helix formation time series, peptides

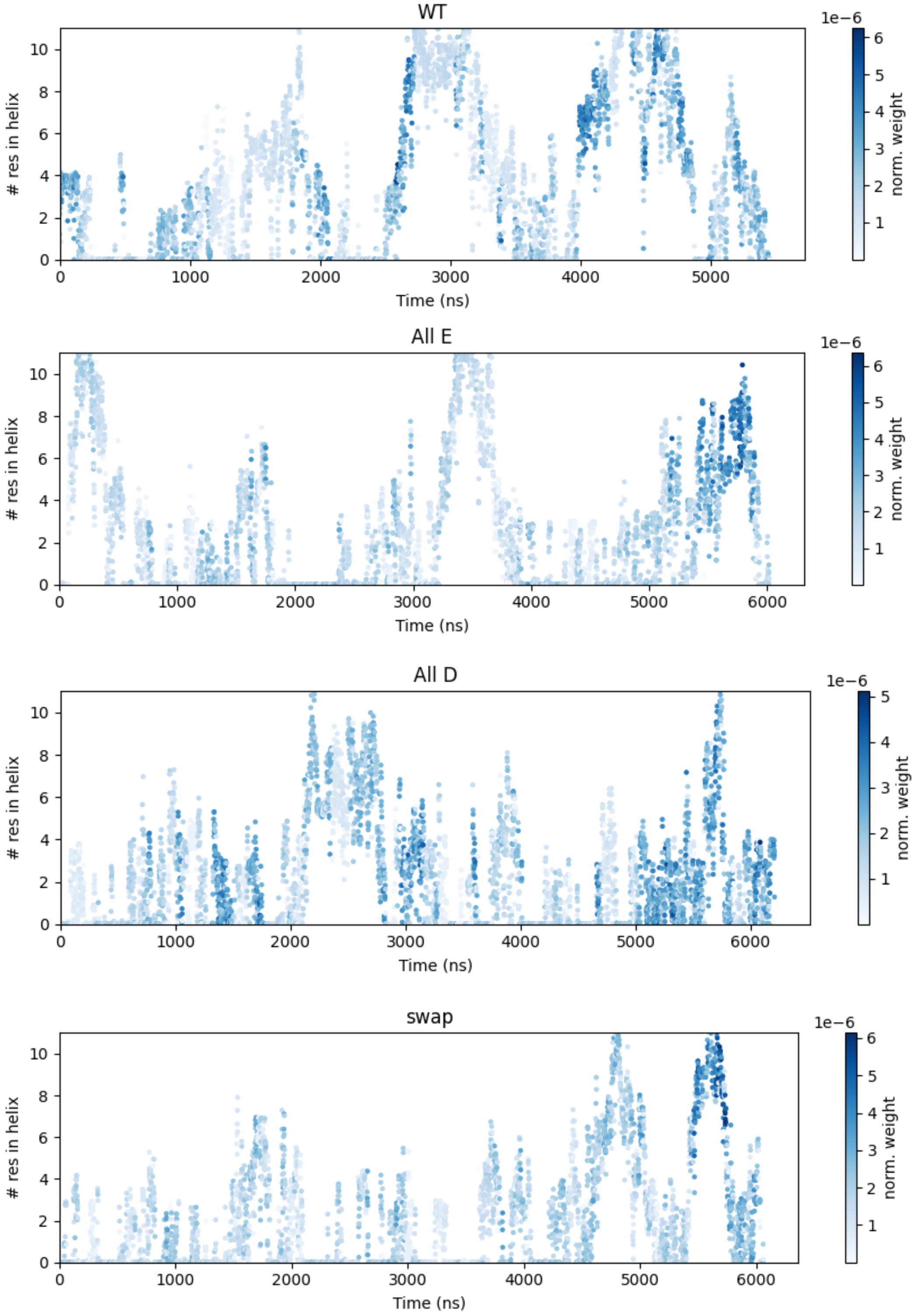


Figure S6: Time series of the number of residues in a helix (α, $3_{10}$ or π) in the simulations of the helix region of Dss1 wildtype and variants, weighted by the bias. Helix content is calculated with DSSP and averaged over 1 ns (100 frames).

### Figure S7: Helix formation time series, full length

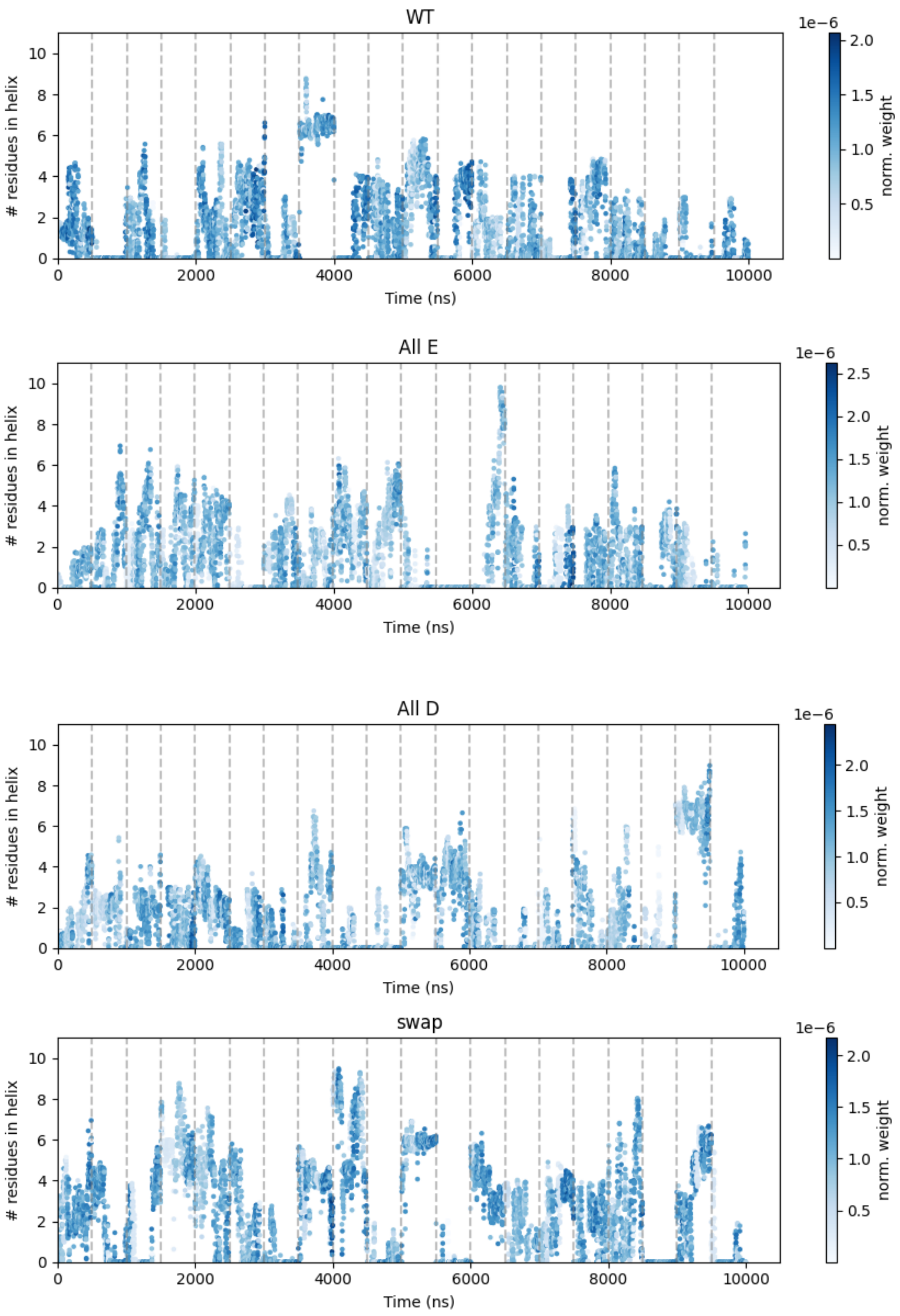


Figure S7: Time series of the number of residues that are in a helix (α, $3_{10}$ or π; residues F55-K66) in the simulation of full-length Dss1 wildtype and variants, weighted by the bias. The replicas are separated by grey dashed lines. Helix content is calculated with DSSP and averaged over 1 ns (100 frames).

### Figure S8: Helix population, simulations

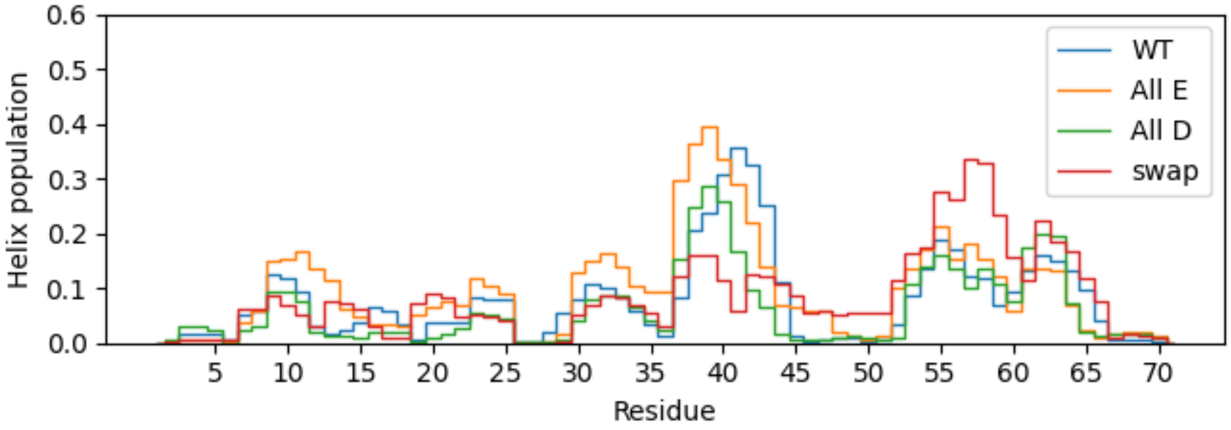


Figure S8: The fraction of the weighted simulation time that each residue is in a helix for full-length Dss1 wildtype and variants, calculated with DSSP. The helix population is shown as the sum of α-, $3_{10}$- and π-helices.

Figure S9: Salt bridge frequency, simulations

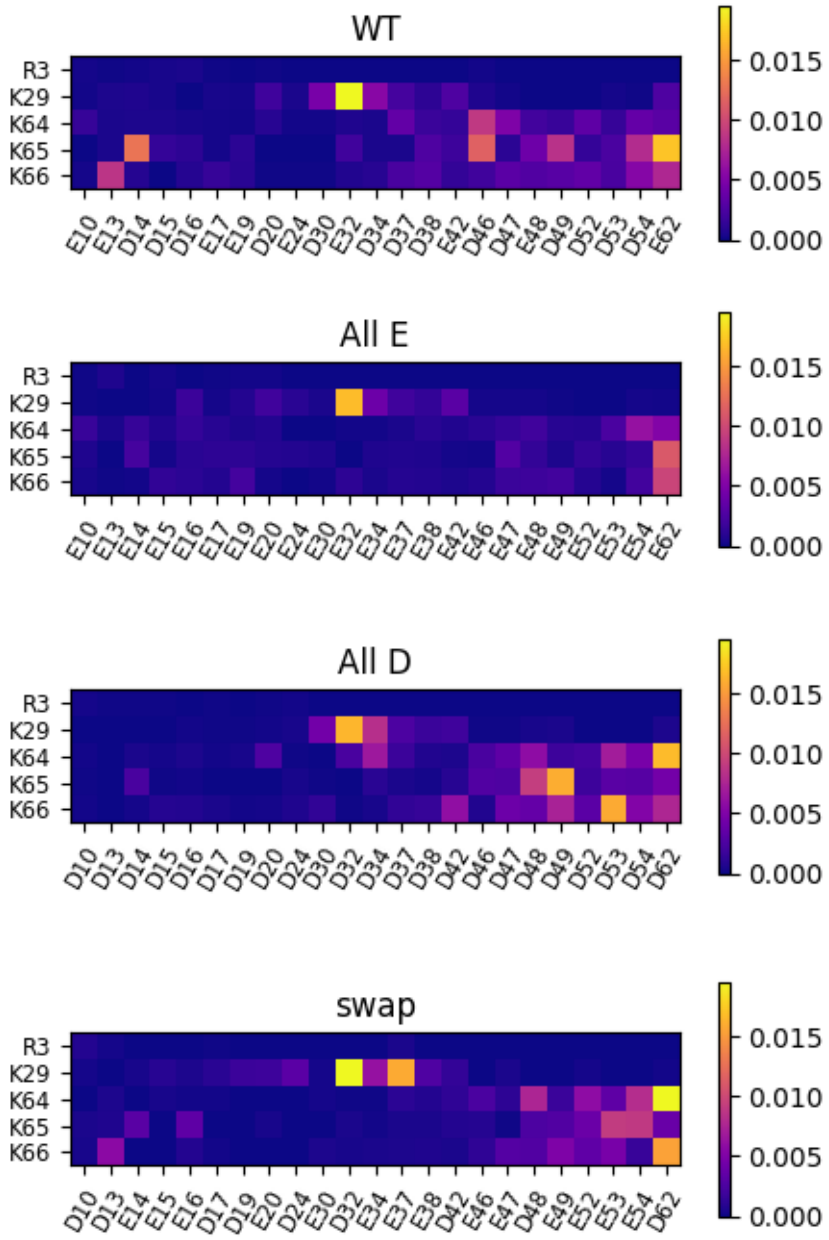


Figure S9: The frequency of each of the possible salt bridges that can be formed in Dss1 relative to the number of frames in each of the full-length simulations of Dss1 and the variants. A salt bridge is defined as a distance of ≤ 3.2 Å between the center of charge of two oppositely charged residues.
